# Analysis of *RDR1/RDR2/RDR6*-independent small RNAs in *Arabidopsis thaliana* improves *MIRNA* annotations and reveals novel siRNA loci

**DOI:** 10.1101/238691

**Authors:** Seth Polydore, Michael J. Axtell

## Abstract

Plant small RNAs regulate key physiological mechanisms through post-transcriptional and transcriptional silencing of gene expression. sRNAs fall into two major categories: those that are reliant on *RNA Dependent RNA Polymerases* (*RDRs*) for biogenesis and those that aren’t. Known *RDR*-dependent sRNAs include phased and repeat-associated short interfering RNAs, while known *RDR*-independent sRNAs are primarily microRNAs and other hairpin-derived sRNAs. In this study, we produced and analyzed small RNA-seq libraries from *rdr1/rdr2/rdr6* triple mutant plants. Only a small fraction of all sRNA loci were *RDR1/RDR2/RDR6*-independent; most of these were microRNA loci or associated with predicted hairpin precursors. We found 58 previously annotated microRNA loci that were reliant on *RDR1*, −*2*, or −*6* function, casting doubt on their classification. We also found 38 *RDR1/2/6*-independent small RNA loci that are not *MIRNAs* or otherwise hairpin-derived, and did not fit into other known paradigms for small RNA biogenesis. These 38 small RNA-producing loci have novel biogenesis mechanisms, and are frequently located in the vicinity of protein-coding genes. Altogether, our analysis suggest that these 38 loci represent one or more new types of small RNAs in *Arabidopsis thaliana*.

**Significance Statement:** Small RNAs regulate gene expression in plants and are produced through a variety of previously-described mechanisms. Here, we examine a set of previously undiscovered small RNA-producing loci that are produced by novel mechanisms.

## Introduction

RNA DEPENDENT RNA POLYMERASES (RDRs) are a family of proteins that produce double-stranded small RNA (sRNA) precursors by synthesizing the complementary strand of a single-stranded RNA molecule (Baulcombe., 2004; Wassenegger and Krczal., 2006; Willmann et al., 2011). Six *RDRs*, divided into the *RDRα* (consisting of *RDR1, RDR2,* and *RDR6*) (Leibman et al. 2017) and *RDRγ* sub-clades (consisting of *RDR3, RDR4*, and *RDR5*) (Willmann et al., 2011), are encoded by the *Arabidopsis thaliana* genome. RDR6 in concert with the RNAse-III like enzyme DICER-LIKE 4 (DCL4) is necessary for the production of phased short interfering RNAs (siRNAs) (Bouché et al., 2006; Howell et al., 2007). RDR2 and DCL3 are involved in the biogenesis of repeat-associated siRNAs (Matzke et al., 2009). RDR1 is involved with defense responses by converting viral-derived single-stranded RNAs into double-stranded RNAs (Leibman et al. 2007), which are then metabolized by DCL4 (or DCL2 when DCL4 function is ablated) (Bouché et al., 2006; Diaz-Pendon et al., 2007). At least one study suggests that *RDR1* is involved in endogenous gene regulation as well (Lam et al., 2012). In contrast to the *RDR*α genes, the functions of the *RDRγ* genes have not yet been described (Willmann et al., 2011).

Not all sRNAs are reliant on *RDRs* for biogenesis: known classes of *RDR*-independent sRNAs include microRNAs (miRNAs) (Vaucheret et al., 2004), which are precisely processed from short (typically less than 300 bases) hairpin RNA molecules (Reinhart et al., 2002), and long inverted repeat siRNAs (long ir-siRNAs) (Henderson et al., 2006), which are imprecisely processed from longer hairpins (the two best described irRNAs, *IR71* and *IR2039,* are more than one thousand nucleotides long) (Henderson et al., 2006). A third class, natural antisense siRNAs (nat-siRNAs), has been reported in various studies (Wang et al., 2005; Jin et al. 2008; Wang et al., 2014). Nat-siRNAs are derived from two distinct, partially complementary transcripts that hybridize to produce an *RDR*-independent double-stranded sRNA precursor (Borsani et al., 2005; Jin et al., 2008; Zhang et al., 2012; Zhang et al., 2013).

Since their discovery the biogenesis of plant sRNAs has been intensely studied, there are yet several unknowns. One open question is whether or not there are any other *RDRα*-independent mechanisms of sRNA biogenesis besides hairpin processing. Another open question is to what extent, if any, the *RDRγ* genes contribute to the sRNA transcriptome. At least one study has shown that the *RDRγ* genes contribute to anti-viral defense in *Solanum chilense* (Verlaan et al., 2013), but how they do so is unknown.

In this study, we produce and utilize small RNA datasets from *rdr1-1/rdr2-1/rdr6-15* (*rdr1/2/6*) triple mutants and use stringent analytical methods to address these questions. Using these data we re-evaluate currently existing small RNA annotations. Ultimately, we identified 38 *RDR*-independent small RNA-producing loci that do not fit into any currently understood schema for plant small RNA biogenesis.

## Results

### Small RNA-seq from *rdr1/2/6*

Small RNAs were sequenced from *Arabidopsis thaliana* wild-type (Col-0) and *rdr1-1/rdr2-1/rdr6-15* triple mutant (hereafter *rdr1/2/6*) mature inflorescences. Three biological replicate libraries of each genotype were obtained. *rdr1/2/6* plants showed a marked decrease in the proportion of 24 nt sRNAs compared to wild-type (Figure 1a), presumably due to loss of *RDR2*-dependent 24 nt siRNAs, which dominate the *A. thaliana* sRNA profile (Kasschau et al., 2007). In contrast, the proportion of 21 nt sRNAs showed a sizeable increase in *rdr1/2/6* datasets compared to wild-type (Figure 1a). This is likely due to the fact that the fraction of sRNAs aligning to *MIRNA* loci is much higher in *rdr1/2/6* libraries than in wild-type libraries (Figure 1b), and most miRNAs in *A. thaliana* are 21 nt in length (Kasschau et al., 2007).

**Figure 1.**
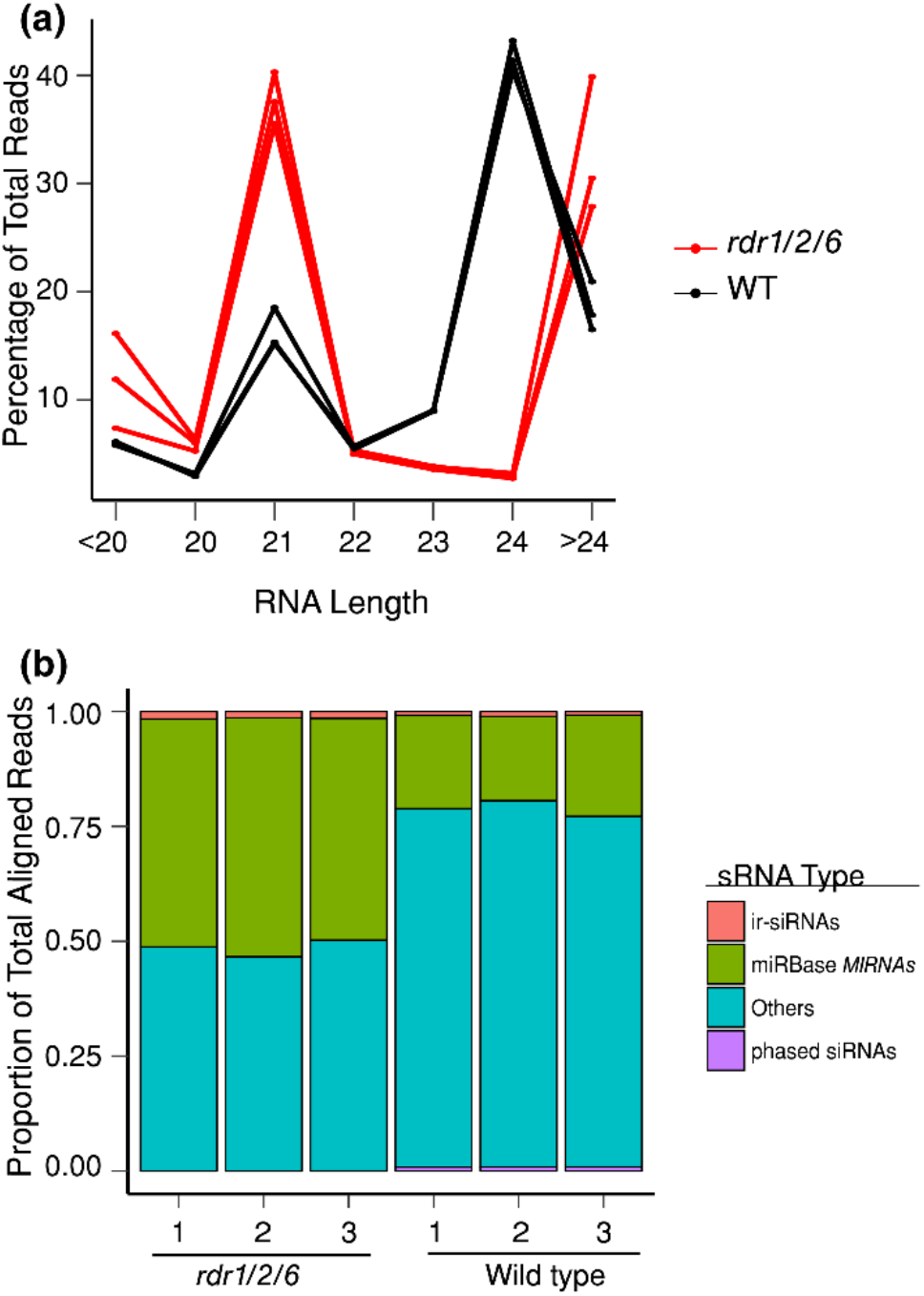
Wild Type and *rdr1/2/6* libraries have distinct sRNA lengths and accumulation patterns. (a) Size distribution of trimmed reads in wild type and *rdr1/2/6* libraries.
(b) Proportion of aligned reads to indicated types of sRNA loci in wild-type and *rdr1/2/6* libraries.

### Critical inspection of *Arabidopsis thaliana MIRNA* annotations in miRBase21

miRBase, the central registry for *MIRNA* loci and mature miRNA sequences (Kozomara and Griffiths-Jones., 2014), is known to contain many questionable miRNA entries (Coruh et al., 2014; Kozomara and Griffiths-Jones., 2014; Taylor et al., 2014). miRBase version 21 lists just 78 out of the 325 *A. thaliana* loci (24%) as ‘high confidence’ annotations. The accumulation of true miRNAs should be *RDR*-independent because, by definition, *MIRNA* primary transcripts are single-stranded hairpins, not the double-stranded RNAs that are produced by RDR proteins (Meyers et al. 2008). Because *RDR*-dependent loci are so numerous, we hypothesized that they could be a major source of false positives in existing *MIRNA* annotations. We thus analyzed the 325 previously annotated*MIRNAs* relative to our data. 58 miRBase annotated *MIRNAs* were significantly (FDR <=0.1) down-regulated in *rdr1/2/6* libraries (Figure 2; Dataset S1). Four of these down regulated loci are siRNAs derived from long inverted repeat regions (Figure S1).

**Figure 2.**
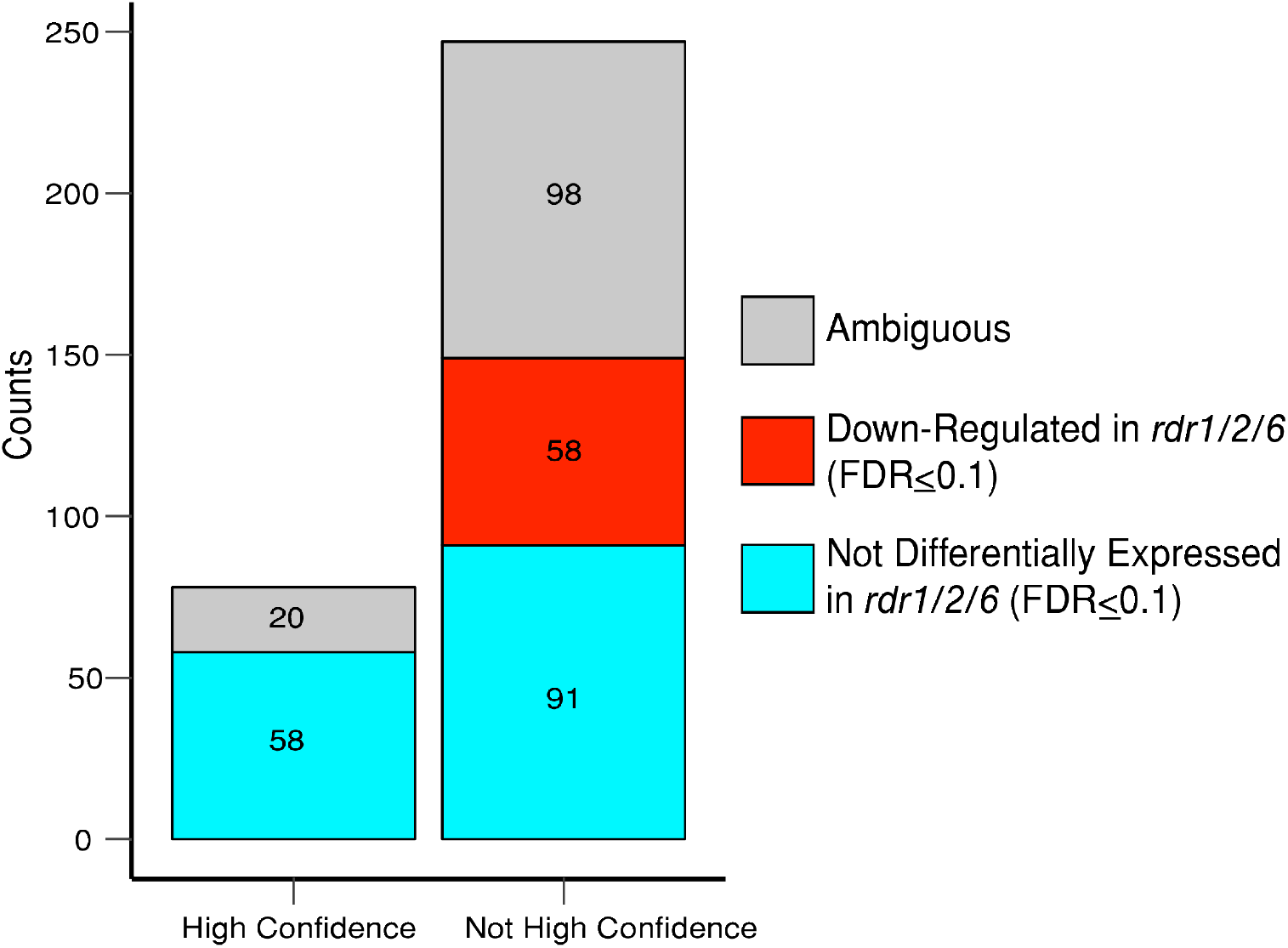
*RDR1/2/6*-dependency of annotated *MIRNAs*. Counts of miRBase21-annotated *MIRNA* loci by “confidence” (per miRBase) and differential expression status. Note that no *MIRNAs* were up-regulated in the *rdr1/2/6* background.

This may have been why some of these siRNA loci were mis-annotated as *MIRNAs.* It also shows that *RDR1/2/6*-dependent siRNAs can overlap with long inverted repeats. However, from these data alone, it’s not clear if the inverted repeat is a DCL substrate used to produce these *RDR1/2/6*-dependent siRNAs or if the siRNA locus is RDR2-dependent and merely overlaps by chance with the inverted repeat region.

The differential expression status of 118 miRBase21 *MIRNAs* was ambiguous because it could not be reliably inferred by our statistical methods (Figure 2). These loci are either too lowly- or too variably-expressed to confidently determine their dependence on *RDRs* based on sRNA-seq data. The remaining 149 miRBase21 *MIRNAs* are not differentially expressed in our libraries and thus are *RDR*-independent (Figure 2), as expected for true *MIRNAs.* This includes 91 *MIRNAs* that aren’t annotated as high confidence loci in miRBase 21 (Figure 2).

### Most small RNA-producing loci are down-regulated in *rdr1/2/6*

We inferred small RNA clusters genome-wide and performed differential expression analysis (Dataset S2; Figure 3a). 33,115 loci either mapped to plastid-derived sequences, did not have reads predominantly in the 20-24 nt range, or had ambiguous differential expression status (Dataset S2). The other 32,859 loci produced reads predominantly in the 20-24 nt range had a differential expression status that could reliably inferred using our statistical tests (Dataset S2; Figure 3a). 32,681 (over 99%) loci were down-regulated in *rdr1/2/6* triple mutants (Figure 3b). The majority of the down-regulated sRNA loci in *rdr1/2/6* are 24 nt siRNA dominated (roughly 98%) (Figure 3b), as expected for a background that contains a null *rdr2* allele (Matzke et al., 2009). 176 loci were found to be confidently not differentially expressed in *rdr1/2/6* triple mutants relative to wild-type (Figure 3a and Figure 3b) by our statistical methods. 112 (~64%) of these loci predominantly produced 21 nt small RNAs (Figure 3b). Only two small RNA loci were up-regulated in the *rdr1/2/6* background (Figure 3a and Figure 3b).

**Figure 3.**
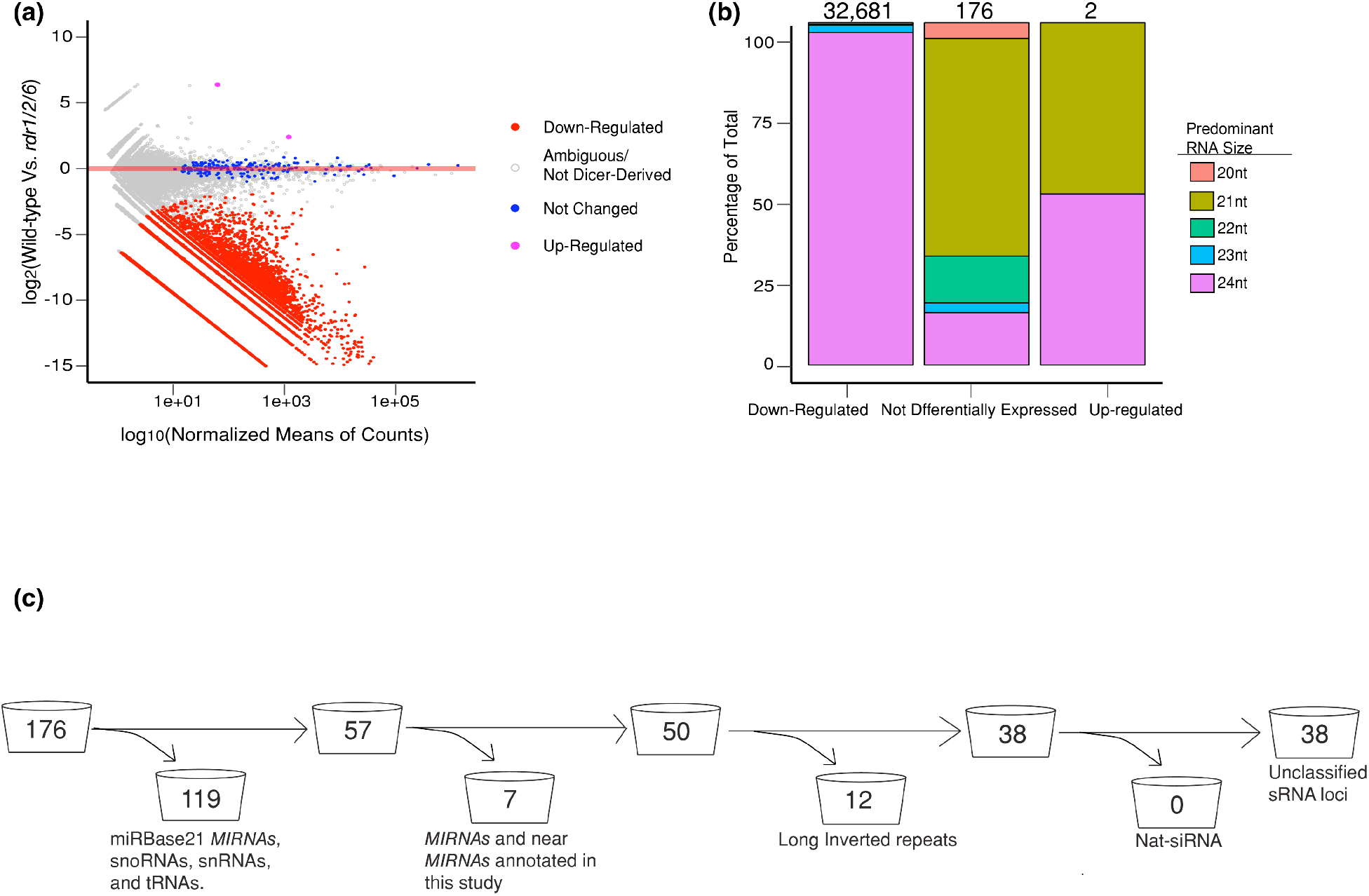
Annotation of *de novo* sRNA clusters. (a) Colored dots represent clusters of distinct differential expression status (FDR<=0.1) based on DESeq2 analysis between wild-type and *rdr1/2/6* mutants.
(b) Counts of loci by differential expression status and predominant RNA sizes.
(c) Analysis pipeline for *RDR1/2/6*-independent sRNA clusters.

As a control, we examined the differential expression status of 41 previously described *RDR6*-dependent loci (Howell et al., 2007) and 11 previously described *RDR2*-dependent loci (Matzke et al., 2009). As expected, 41 out of 52 of the loci were down-regulated in *rdr1/2/6* triple mutants (Dataset S3). The differential expression status of the other 11 known *RDR6*-dependent loci couldn’t be reliably determined through our statistical methods, due to low read counts and/or variability in the replicates.

Of the 176 small RNA loci that weren’t differentially expressed, 119 (roughly 68%) intersected with annotated*MIRNAs,* tRNAs, rRNAs, or snoRNAs (Figure 3c). This left 57 loci, two of which we annotated as novel *MIRNAs* (Figure 3c; Figure S2a; Dataset S2). We classified another five loci as MIRNA-like (Figure 3c; Dataset S2). These were either hairpin loci in which the predicted miRNA* either wasn’t sequenced or the miRNA/miRNA* sequences were imprecisely processed (Figure S2b), or loci that were less than 400 nts and had at least 90% overlap with predicted hairpins. Another 12 loci were annotated as long inverted repeats (ir-siRNAs) (Figure 3c), including the previously described locus *IR71* (Henderson et al., 2006; Dataset S4). ir-siRNAs were those loci in which the small RNAs were imprecisely processed from one or more regions with predicted stem-and-loop secondary structure (Figure S2c). We did not find any nat-siRNAs among the *RDR1/2/6*-independent loci (Figure 3c). The remaining 38 *RDR1/2/6*-independent loci were not placed into any currently known paradigms of plant sRNA production (Dataset S4; Figure S2d).

### Unclassified *RDR1/2/6*-independent loci are true small RNA-producing clusters that are 21 nt or 24 nt dominated

We hypothesized that inaccurate placement of multi-mapping reads at erroneous locations could have potentially contributed to the apparent small RNA production at the 38 unclassified *RDR1/2/6*-independent loci. We therefore examined the percentage of multi-mapping reads at all of the sRNA loci. This hypothesis was not strongly supported as the unclassified *RDR1/2/6*-independent loci produced very few multi-mapping reads (Figure 4a), indicating that these reads were generally not derived from other loci. Generally, the unclassified *RDR1/2/6*-independent loci show modest-to-high expression levels compared with the other classes of loci (Figure 4b). This further indicates that these loci are true small RNA-producing loci instead of just alignment artifacts.

**Figure 4.**
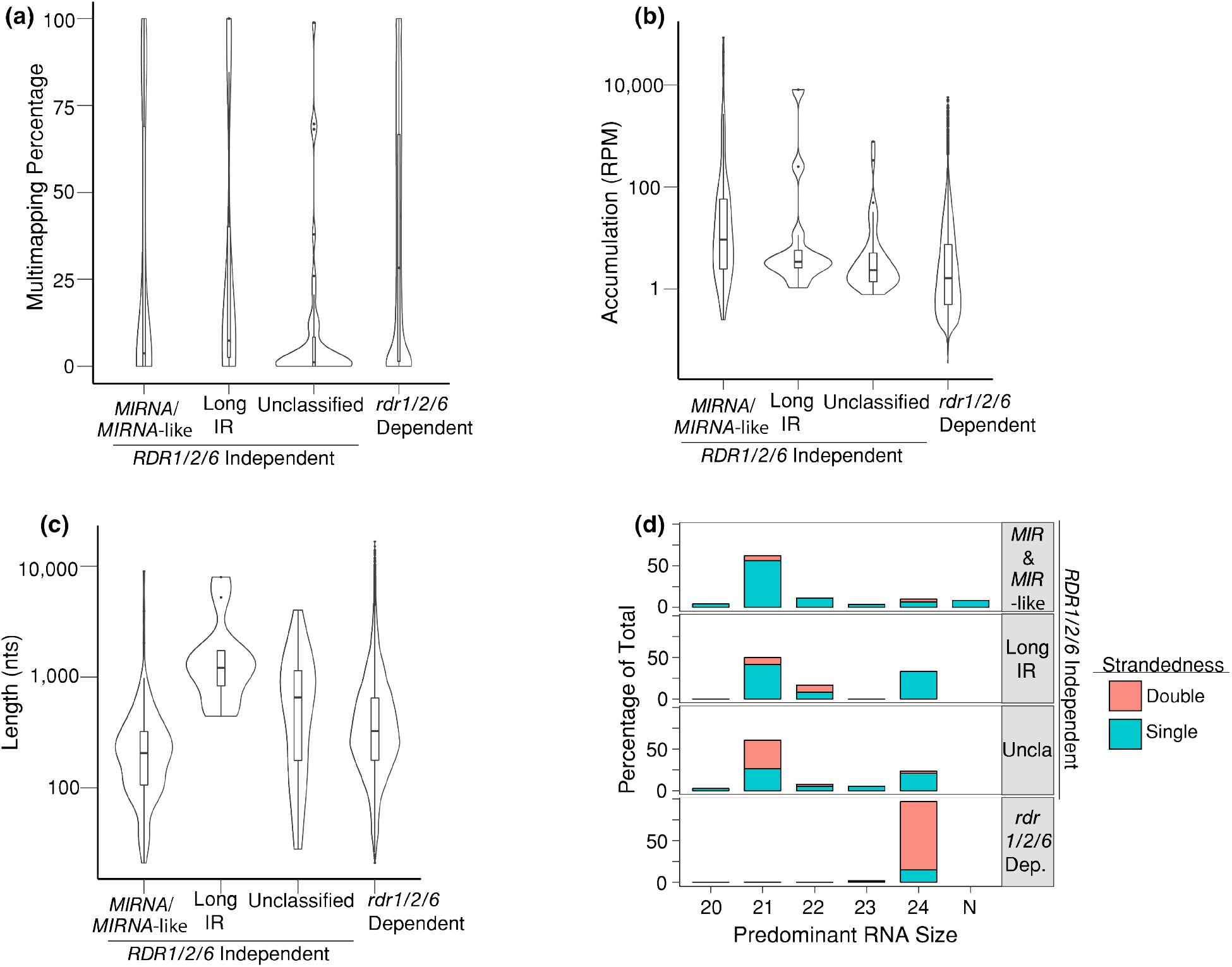
Unclassified loci are true sRNA-generating loci. (a) Percentages of multimapping reads in wild-type libraries for different types of loci. The width of the density plot shows the frequency. The inset boxes show medians (horizontal lines), the 1st-3rd quartile range (boxes), the 95% confidence of medians (notches), other data out to 1.5 times the interquartile range (whiskers) and outliers (dots).
(b) Same as panel (a) except for sRNA accumulation (RPM: Reads per Million)
(c) Same as panel (a) except for locus lengths.
(d) Percentage of predominant RNA size and strandedness for different types of loci from wild-type libraries.

Unclassified *RDR1/2/6*-independent loci had a heterogenous size distribution, with most clusters falling between 100 and 1,000 nts (Figure 4c). We were interested in determining whether or not these loci were synthesized by different sRNA biogenesis mechanisms. To test this hypothesis, we inferred the expression of these clusters in various mutants related to sRNA biogenesis using previously published datasets (specific genotypes and datasets are listed in Table S1). Most of the unclassified *RDR1/2/6*-independent loci are either ambiguous or not differentially expressed in the backgrounds tested (Figure S3).

Most unclassified *RDR1/2/6*-independent loci predominantly produce 21 nt or 24 nt small RNAs (Figure 4d). Many of the unclassified, 21 nt *RDR1/2/6*-independent loci were doublestranded (sRNA reads derived from both polarities of the genome; Figure 4d). This suggests processing from double-stranded precursors. This is of interest because these are double stranded RNA loci produced independently of the biochemical activities of RDR1, RDR2, and RDR6. It is worth noting that long ir-siRNAs loci are also overwhelmingly double stranded in origin (Figure 4d). This could be due to their near perfect palindromic hairpins precursors leading to ambiguously mapped reads on both arms of the IR (Zhang et al., 2006). This does not, however, explain the double-stranded nature of these unclassified *RDR1/2/6*-independent loci since they do not overlap detectable long inverted repeats or hairpins. These results suggest that these unclassified *RDR1/2/6*-independent loci are derived from potentially novel small RNA-producing mechanisms.

### Unclassified loci overlap with genes but are not produced from their transcripts

Most (76%; 29 of 38 clusters) of the unclassified *RDR1/2/6*-independent small RNAs loci overlapped with genes (Figure 5a; Dataset S5). This proportion of genic overlap was higher than the other categories of small RNAs loci described in this study (Figure 5a). In contrast, *rdr1/2/6*-dependent are overwhelmingly transposon-overlapping (~78%) (Figure 5a), likely because of the prevalence of *rdr2*-dependent loci. We were interested in determining why the unclassified loci would localize to genes. One trivial explanation is that the small RNAs at these overlapping loci were simply degraded fragments of mRNA transcripts where by chance at least 80% of the reads happened to be between 20 and 24 nts long. We therefore placed the gene-intersecting unclassified *RDR1/2/6*-independent clusters into three different categories based on their genomic polarities relative to the genes they overlapped (Figure 5b). Same-stranded clusters are consistent with degraded transcripts (Figure S4a), while opposite-stranded (Figure S4b) and double-stranded clusters (Figure S4c) are not. Of the 29 gene-overlapping unclassified *RDR1/2/6*-independent loci, only 11 (~38%) were on the same strand (Figure 5b). Therefore, the pattern of small RNA accumulation at more than half of these loci argues against the degraded transcript hypothesis.

**Figure 5.**
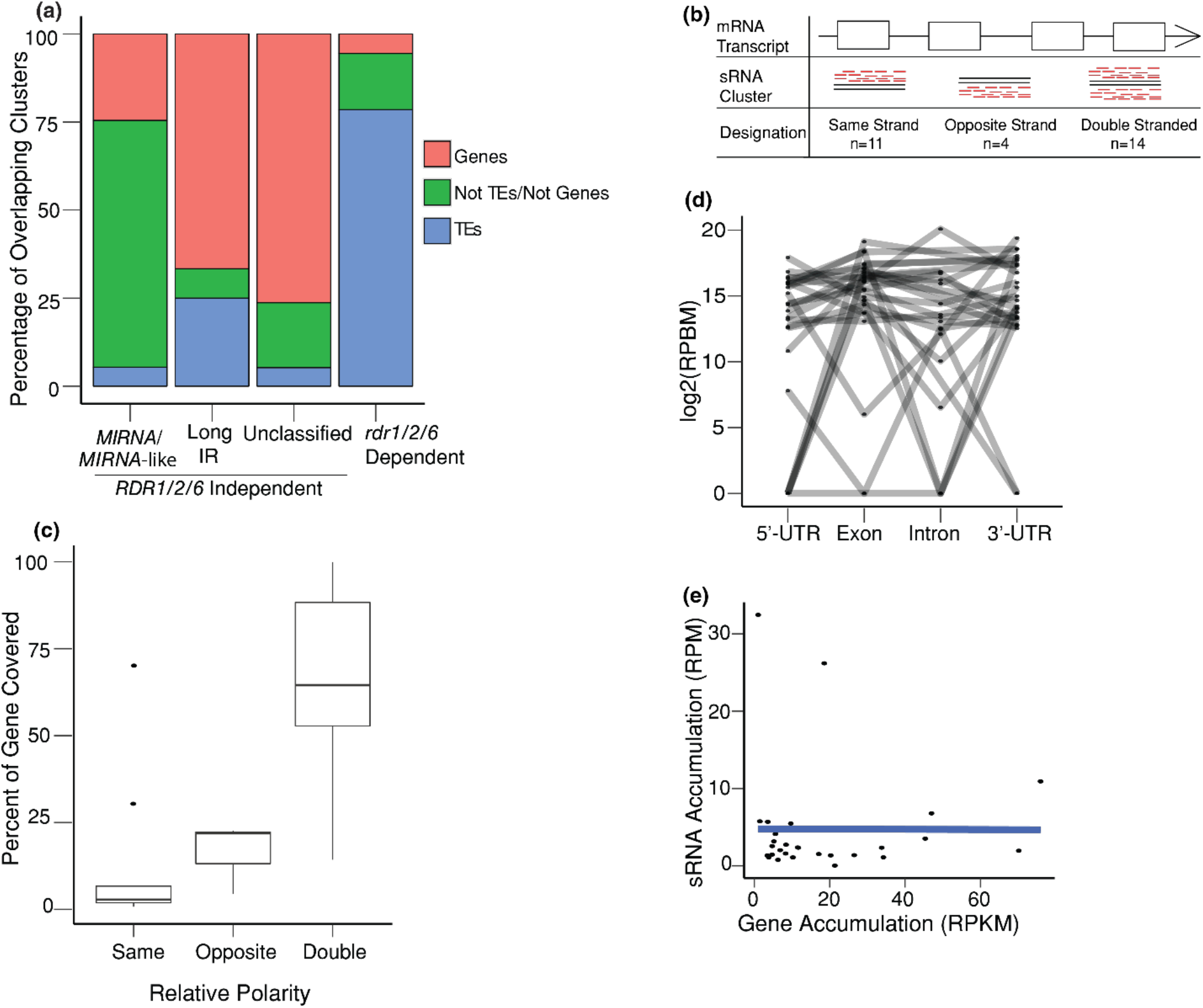
Gene intersecting sRNA clusters are not mRNA fragments. (a) Percentage of unclassified *RDR1/2/6*-independent clusters either intersecting genes or transposable elements, calculated as: (number of clusters intersecting a feature/total number of clusters in category)*100.
(b) Gene intersecting unclassified *RDR1/2/6*-independent sRNA loci were designated based on their genomic polarities relative to the genes they intersected.
(c) Percentage of genes covered by intersecting unclassified *RDR1/2/6*-independent sRNA cluster(s).
(d) Number of reads mapped to different portions of unclassified *RDR1/2/6*-independent sRNA-intersecting genes. (RPBM: Reads per Base Millions).
(e) Correlation of the accumulations of genes and their overlapping sRNA clusters. (RPKM: Reads per Kilobase Million and RPM: Reads per Million).

We next tested the amount of overlap between the same-stranded loci and different portions of the overlapping gene in order to determine if the same-stranded loci were simply transcript fragments. If the small RNAs are simply transcript fragments, we would expect the originating cluster to completely cover the gene. Instead, most of the same-stranded small RNA loci cover less than 25% of their overlapping gene (Figure 5c), suggesting that the small RNAs at these same stranded clusters are not simply mRNA transcript fragments and that there may even be “hotspots” in the genes more conducive to DCL processing. This could suggest that the mature mRNA transcript or at least portions of it are utilized by DCLs to produce sRNAs at these loci. In this case, we would expect significantly less reads mapping to introns, as introns are not a part of the mature mRNAs, and a strong correlation between the expression of the unclassified *RDR1/2/6*-independent loci and their overlapping genes. Only two of the genes had no aligned sRNAs within introns (Figure 5d). The other genes either had reads mapped to introns or lacked introns altogether (Figure 5d). Furthermore, there is not a strong positive correlation the expression of overlapping unclassified *RDR1/2/6*-independent cluster and their intersected genes (Figure 5e). Altogether our results suggest that the majority of unclassified *RDR1/2/6*-independent loci localize to certain genes but are not produced directly from their mRNAs.

### The unclassified loci are largely not conserved

Why do many of the unclassified small RNA clusters overlap with genes if the mRNAs aren’t used as DCL substrates? A simple explanation is that these are evolutionarily young small RNA loci which are reliant on the transcriptional machinery that localizes to the genes in order to be transcribed. Indeed, it has been reported that new genes are typically born around or even on the opposite strand of previously existing genes (Siepel, 2009; Li et al. 2010; Guerzoni and McLysaght, 2016). We therefore investigated the sequence identity of these small RNA loci between nine species in the Brassicaceae (Haudry et al., 2013) and quantified the conservation levels (using an arbitrary unit of measurement). We found that most of the unclassified *RDR1/2/6*-independent loci are not detectably conserved with a median conservation score of 1, though some are moderately conserved when compared to the other small RNA clusters analyzed in this study (Figure 6). We also note similar results for all other types of small RNA loci described in this study except for *MIRNA/MIRNA-like* loci, some of which are highly conserved (Zhang et al., 2006; Fahlgren et al., 2007).

**Figure 6.**
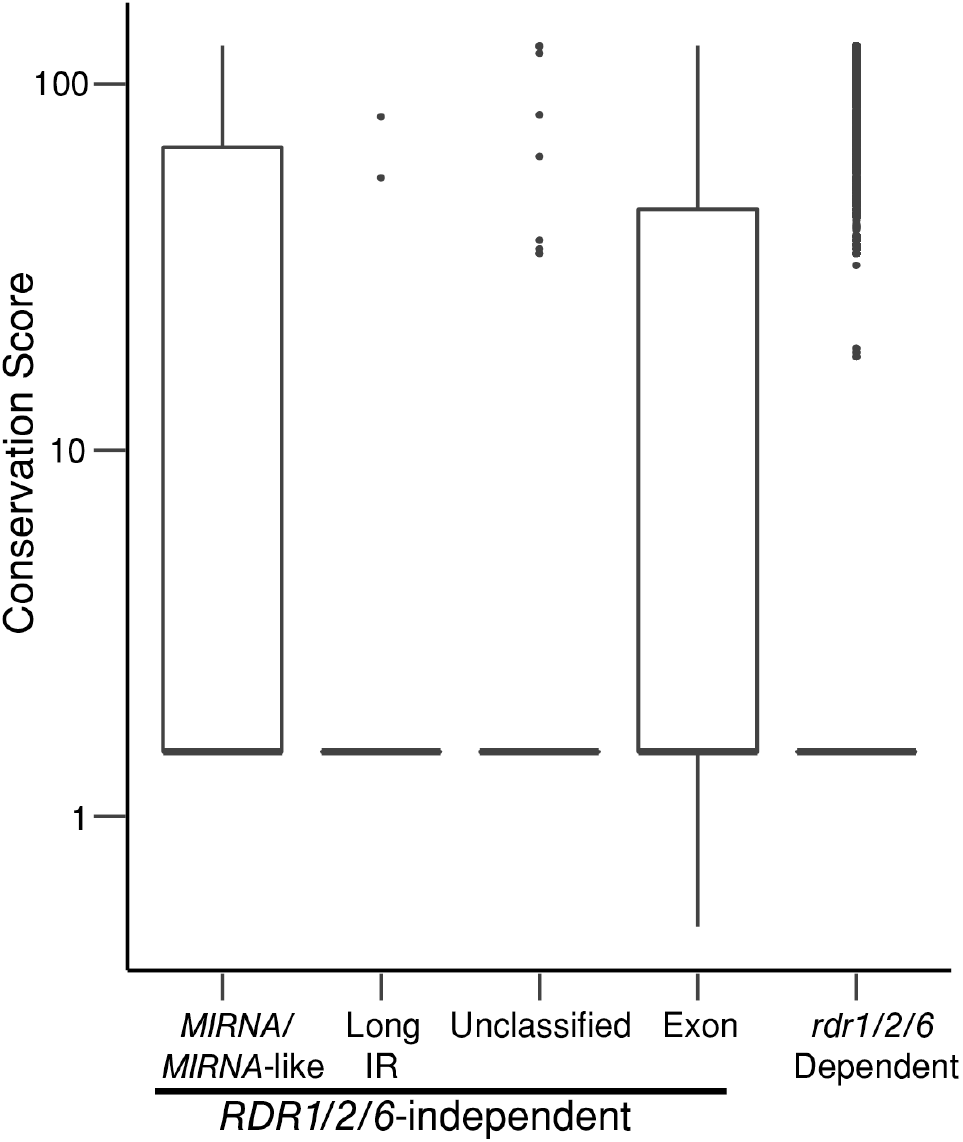
Conservation analysis of sRNA loci. Conservation scores were assigned to each read in the *A. thaliana* (TAIR10) genome based on whole-genome alignments between nine Brassicaceae species. Boxplots representing the median conservation scores of indicated types of sRNA loci are shown.

### The *rdr6-15* T-DNA insertion causes sRNA production at the *RDR6* locus

Only two small RNA loci were up-regulated in the *rdr1/2/6* triple mutant (Figure 1a). One up-regulated locus overlaps with the *RDR6* gene, particularly at the T-DNA insertion site in the *rdr6-15* allele (Figure 7a). We conclude that the T-DNA insertion at the *rdr6-15* allele has triggered sRNA biogenesis at the *RDR6* locus. However, no small RNAs were detected at the T-DNA insertion sites of *rdr2-1* (Figure 7b) or *rdr1-1* (Figure 7c).

**Figure 7.**
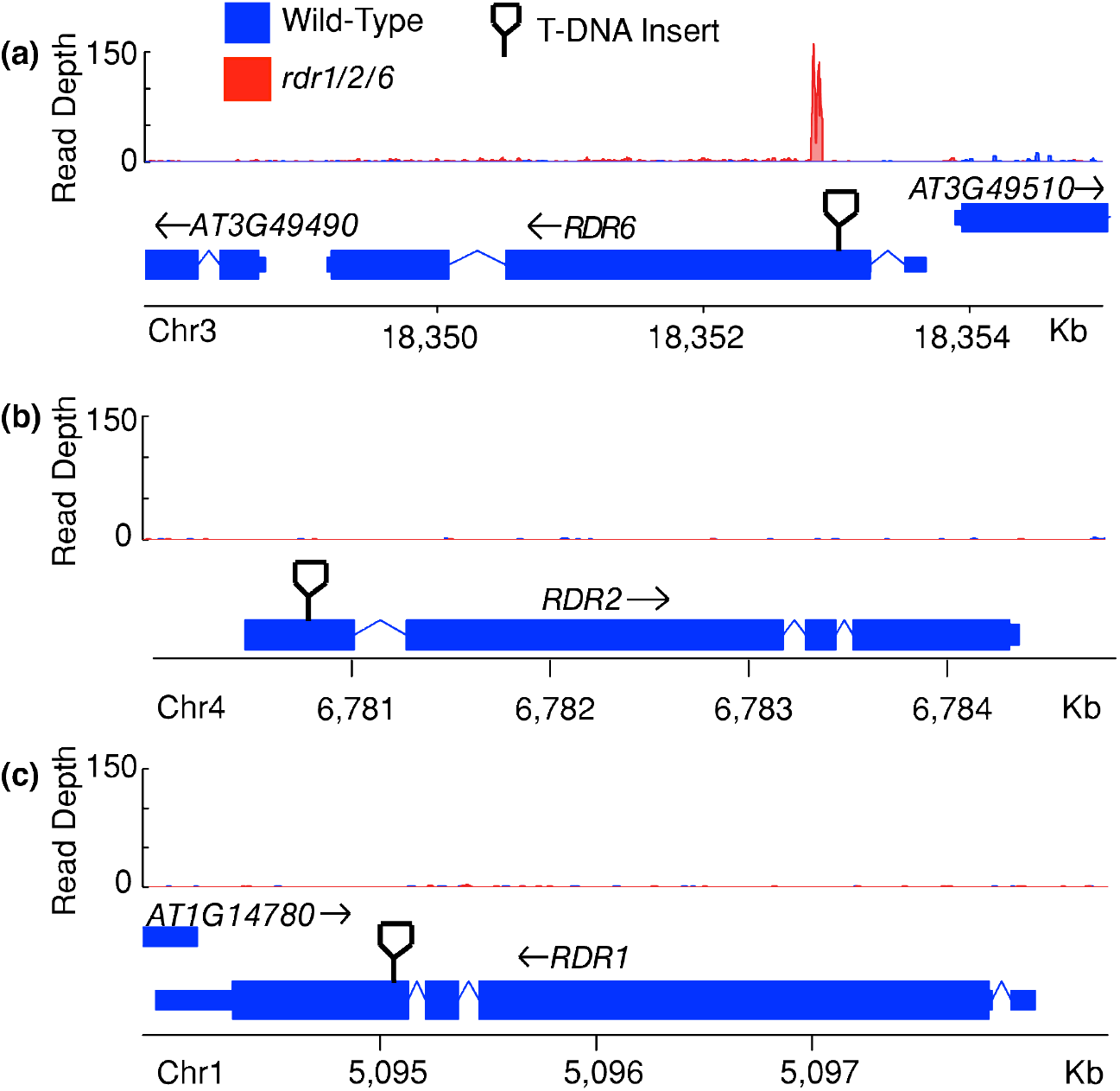
*RDR6* overlaps with an up-regulated sRNA cluster. (a) Histogram of read depth for an up-regulated unclassified sRNA locus derived from the T-DNA insertion site of *rdr6-1*. Arrows represent genomic polarity of locus. Two arrows mean that the particular locus is derived from both strands.
(b) Histogram of read depth of the *RDR2* locus, showing the T-DNA insertion site of the *rdr2-1* allele.
(c) Histogram of read depth of the *RDR1* locus, showing the T-DNA insertion site of the *rdr1-1* allele.

## Discussion

### Nearly 20% of annotated *A. thaliana MIRNAs* are dependent on *RDRs*

Many *MIRNA* annotations in miRBase are suspected to be erroneous (Coruh et al., 2014; Kozomara and Griffiths-Jones., 2014; Taylor et al., 2014). As *MIRNA* primary transcripts are single-stranded, mature miRNA production is by definition *RNA DEPENDENT RNA POLYMERASE* (RDR)-independent. However, we found 58 annotated *MIRNAs* to be *RDR*-dependent (Figure 2; Dataset S1) and thus we propose that they aren’t true *MIRNAs.* Interestingly, the predicted secondary structure of these erroneous *MIRNAs* were predicted to be hairpin structures (Figure S1), albeit more akin to long inverted repeats rather than the short hairpins that produce miRNAs (Henderson et al., 2006). This opens the possibility of a class of *RDR*-dependent ir-siRNAs, though it’s entirely possible that these long inverted repeats simply happen to overlap *RDR*-dependent loci. In either case, two things are apparent: (i) simply using the secondary structure of sRNA cluster is inefficient for robust sRNA annotation and (ii) more study is necessary to ascertain what the DCL substrate is in these cases. In the future, more tests will be necessary to verify other miRBase annotations.

### Many *RDR1/2/6*-independent sRNA loci are *MIRNAs*, *MIRNA-like*, or hairpin-derived

We had expected that nat-siRNAs would be among the *RDR*-independent small RNA loci, but we found none (Figure 3c). Nat-siRNAs are derived from partially complementary transcripts that serve as the double-stranded sRNA precursors (Jin et al., 2008; Zhang et al., 2012). Zhang et al. (2012) considered any sRNA clusters convergent gene pairs to be nat-siRNAs, even if they weren’t necessarily derived from the overlapping portion of the genes. However, we only considered a small RNA cluster to be nat-siRNA-producing if it intersected the overlapping portion of the convergent gene pairs: no *RDR*-independent sRNA cluster met this criterion. Furthermore, we did not consider the *rdr1/2/6*-dependent sRNA clusters in our nat-siRNA annotation because, by definition, the biogenesis of nat-siRNAs should be *RDR*-independent. One possibility is that nat-siRNAs simply aren’t produced in the conditions examined in this study (wild-type inflorescences in the absence of biotic and abiotic stresses). Borsani et al. (2005), and Wang et al. (2014) described a nat-siRNA in salt stress conditions and light stimulus, respectively.

Among the classified *RDR1/2/6*-independent loci, we describe several MIRNA-like loci (Figure 3c). While some of these may be true *MIRNAs* in which the miRNA* simply wasn’t sequenced, we note that many MIRNA-like loci imprecisely produce reads (Figure S2b). Previous studies have hypothesized that are long ir-siRNAs are actually *proto-MIRNAs* (Allen et al., 2004; Fahlgren et al., 2007). Curiously, these MIRNA-like loci show properties of both *MIRNAs* (produced from shorter hairpins) and long ir-siRNAs (producing constellations of different sizes of sRNA reads; Figure S2b). These MIRNA-like loci could represent transitional forms between long ir-siRNAs and *MIRNAs.* However, future studies to investigate this hypothesis might be impeded by the low conservation of these clusters (Figure 7). This will make cross-species analyses difficult.

### Unclassified, *RDR1/2/6*-independent sRNA loci represent potentially novel regulatory RNAs

38 *RDR1/2/6*-independent sRNA clusters couldn’t be classified indicating that they could be made via potentially novel biogenesis mechanisms (Figure 3c). Most of these unclassified sRNA clusters localize to genes (Figure 4d), although the mature mRNAs of these genes are generally not utilized to produce the sRNAs (Figure 5b). It’s possible that these unclassified loci are newly emergent sRNA clusters that are born around other genes; new genes are typically born around already existing genes at least in humans (Siepel, 2009; Li et al. 2010; Guerzoni and McLysaght, 2016) and it’s possible that the same occurs with sRNA clusters in *A. thaliana.* Although many of these unclassified loci are more highly expressed than *rdr1/2/6*-dependent sRNAs, they are typically more lowly expressed than most*MIRNAs* (Figure 4b). It’s possible that many of these unclassified sRNAs are too lowly expressed to be functional.

This study is not the first to describe gene-derived sRNAs in plants. Many phasiRNAs are genic (Howell et al., 2007; Fei et al., 2013). Furthermore, there are many studies that show that in certain genetic backgrounds or viral infections, many mRNAs can be used as sRNA precursors via *RDR* activity (Gazzani et al., 2004; Belostotsky and Sieburth, 2009; Zhang et al., 2015). However, there are key differences between the unclassified, *RDR1/2/6*-independent sRNA clusters and previously described gene-derived sRNA loci. First, previously described gene-derived sRNAs are processed from mRNAs as opposed to merely overlapping the gene, as is the case with these unclassified, *RDR1/2/6*-independent sRNA loci (Figure 5b-c). Furthermore, previously described gene-derived sRNAs are made from transcripts of defense response, pentatricopeptide repeat, and transcription factor genes (Howell et al., 2007; Gregory et al., 2008). The genes overlapping these unclassified, *RDR1/2/6*-independent sRNAs are not obviously enriched in any particular gene ontology, though several pentatricopeptide repeat and defense response related genes are represented (Dataset S5).

Perhaps most interesting is the rather low number of *RDR1/2/6*-independent sRNA clusters. The majority of the *RDR1/2/6*-independent loci were *MIRNAs* or other loci associated with predicted hairpin RNAs. *RDRs* are divided into two clades but only one clade (*RDRα*: *RDR1*, *RDR2*, and *RDR6*) is currently known to be involved in sRNA biogenesis (Willmann et al. 2011). It’s possible that one or more of the *RDRγ* members (*RDR3*, *RDR4*, and *RDR5*) are required for the biogenesis of the 38 unclassified, *RDR1/2/6*-independent sRNA loci that we’ve found. The low number of *RDR1/2/6*-independent sRNA clusters imply that the *RDRγ* genes contribute minimally, if at all, to the endogenous sRNA populations of healthy inflorescence tissues. Further work will be required to fully understand the biogenesis of the unclassified, *RDR1/2/6*-independent sRNA clusters.

## Experimental procedures

### Plant materials and growth conditions

*rdr1-1/rdr2-1/rdr6-15* (*rdr1/2/6*) triple mutant seed was obtained from The Arabidopsis Biological Resource Center (stock number CS66485). All plants were grown at 24° C in 16 hours day/8 hours night lighting conditions. Every 10.5 liters of soil was supplemented with 25ml of Osmocote®.

### RNA Extraction and small RNA library construction and alignment

Mature inflorescence tissues were collected from *rdr1/2/6* and Col-0 plants. Total RNA was extracted using Tri-Reagent (Sigma, St. Louis, MO, USA) following the manufacturer’s protocol. Small RNA libraries were prepared using the TruSeq Small RNA Library Preparation Kit (Illumina) following the manufacturer’s protocol. Biological triplicate libraries were sequenced via Illumina HiSeq 2500 at the Genomic Core Facility at the Pennsylvania State University. Raw data has been deposited to the National Center for Biotechnology Information Gene Expression Omnibus under the accession number GSE105262. Library details and accession numbers are given in Table S1.

Small RNA-seq reads from merged triplicate wild-type and *rdr1/2/6* libraries were aligned against the *A. thaliana* genome (version TAIR10), and used to infer small RNA-producing cluster using ShortStack 3.2 (Johnson et al., 2016) with options --pad 75 and --adapter TGGAATTC. Briefly, small RNA generating loci (designated as “Clusters” in all proceeding tables) were defined as regions in which at least two unique small RNA reads are 300 nts or less apart and at least 5 reads map to the region (Dataset S2).

### Differential expression analysis

Mature high confidence *MIRNA* hairpins sequences for *A. thaliana* were downloaded from miRBase (version 21) (Kozomara and Griffiths-Jones, 2014). The overall proportions of mature miRNAs in each library was then calculated by dividing the total quantity of mature high confidence miRNA sequence in each library by the total number of reads in that library. Read counts of the library were normalized by dividing by the median of the proportion of mature high confidence miRNA among all the libraries. The log2 transformed fold change was then calculated using DeSeq2 (Love et al. 2014). Next, the raw read counts in *rdr1/2/6* libraries were normalized using the median of the fold changes of clusters intersecting high confidence *MIRNAs.* Three differential expression tests were then performed using DeSeq2 (Love et al. 2014). (i) Up-regulated loci were determined at a false discovery rate of 0.1 with an alternative hypothesis that the log2 transformed fold change was greater than 1.5. (ii) Down-regulated loci were determined at a false discovery rate of 0.1 with an alternative hypothesis that the log2 transformed fold change was less than −1.5. (iii) Finally, loci that were not differentially expressed were determined at a false discovery rate of 0.1 with an alternative hypothesis that the log2 transformed fold change was between −1.5 and 1.5. Loci where the null hypothesis was not rejected in all three tests were classified as ambiguous with respect to *RDR1/2/6* dependency (Dataset S2).

### Inspection of miRBase annotated *MIRNA*

Known *MIRNA* loci for *A. thaliana* were downloaded from miRBase version 21 (Kozomara and Griffiths-Jones, 2014). Wild-type and *rdr1/2/6* small RNA libraries were aligned to the *A. thaliana* genome as in our previous differential expression analysis, except a file containing the *MIRNA* loci from miRBase was used as a “locifile” in the input to ShortStack. Furthermore, we used ShortStack version 3.8.1 for these alignments instead of the previously used ShortStack version 3.2; the alignment parameters, however, are similar between both versions. Read count normalization and differential expression were performed as before.

### Designation of small RNA types

Small RNA loci designated by ShortStack with a ‘DicerCall’ of ‘N’, which have less than 80% of their aligned reads between 20 nts and 24 nts in length (inclusive), were removed from further analysis as these tend to be loci that don’t produce regulatory RNA. Loci intersecting the mirBase21 miRNA generating loci, as determined by IntersectBed from the Bedtools version 2.2 suite (Quinlan, 2014), were also removed from further analysis. Finally, the list of regions producing snRNA, snoRNA, and tRNA was downloaded from the EnsemblPlants database (ftp://ftp.ensemblgenomes.org/pub/release-24/plants/gtf/arabidopsis_thaliana/). Loci intersecting these regions as determined by IntersectBed were removed from further analysis.

Novel *RDR*-independent loci were placed into five different categories. (i) *De novo MIRNAs* were those loci that were annotated as such by ShortStack 3.2. (ii) *MIRNA*-like were determined by two methods: (1) annotation by ShortStack. In these cases, these were loci that had imprecise processing of the stem-and-loop secondary RNA structure or were loci in which the miRNA* wasn’t sequenced. (2) An in-house Perl wrapper for RNALFold 2.3.5 (from Vienna RNA Secondary Structure Package version 2.3.5) (Lorenz et al., 2011) was used to find potential hairpins in the *A. thaliana* TAIR10 genome that were at most 400 bps in length and had no more than one-third of their bases unpaired in the predicted secondary structure. These parameters were determined by examining the secondary structures of high confidence *MIRNA* loci. Using this wrapper, both strands of the genome were examined for hairpins. MIRNA-like were those loci that were at least 90% covered by the hairpin loci and had the same strandedness as the hairpin (using IntersectBed with options −f 0.9 and -s), but were less than 400 nts in length. (iii) A list of accession numbers of convergently and divergently overlapping genetic loci as well as genes that were completely enclosed in another gene was downloaded from the Plant Natural Anti-sense Database (Chen et al., 2011). The list of the *A. thaliana* gene models was downloaded from The Arabidopsis Information Resource (ver. TAIR10). The coordinates of the overlapping genes were copied from the TAIR10 gene models and the overlapping proportion of each gene pair was determined via IntersectBed (Quinlan, 2014). Natural antisense siRNA (nat-siRNA) loci were those small RNAs that intersected with the overlapping regions of each gene pair as illuminated by IntersectBed. (iv) Inverted repeats were determined by two different methods. (1) Inverted Repeat Finder ver. 3.07 (Warburton et al., 2004) was used against the *A. thaliana* TAIR10 genome using the following parameters: of a matching weight of 2, a mismatch weight of 3, an indel penalty of 5, a matching probability of 80, an indel probability of 10, and finally GT pairing was allowed with a weighting score of 2. Potential inverted repeat regions in which the loop was longer than the stems were removed. Inverted repeat derived siRNAs (ir-siRNAs) were loci that intersected with the stems of the remaining potential inverted repeats. (2) Loci that were at least 90% covered and had the same strandedness as RNALFold-predicted hairpin (using IntersectBed with options −f 0.9 and -s) but were over 400 nts in length. (v) Finally, unclassified loci were those clusters that simply couldn’t be put into any of the above categories.

### Determining the statistics of different types of loci

Lengths, strandedness, and predominant read size in the locus (“DicerCall” in supplementary tables) were all determined by ShortStack alignment and analysis of merged wild-type libraries against the *A. thaliana* (TAIR10) genome. The results file from our initial ShortStack analysis was used as the locifile. The percentage of multimappers in merged wild-type libraries was calculated by the equation: (1-(number of unique reads at locus/total reads at locus)) * 100.

### Determining overlap between small RNA loci and genetic loci

*A. thaliana* genes (version TAIR10) was downloaded as above and list of transposable elements was downloaded from EnsemblPlants database (ftp://ftp.ensemblgenomes.org/pub/release-24/plants/gtf/arabidopsis_thaliana/). Overlap between these genetic features and the different small RNA loci was determined via IntersectBed (Quinlan, 2014). If a small RNA locus overlapped with both a transposon and a gene, we took into account the ontology of the gene to see if it was a transposable element. Otherwise, it was considered gene-intersecting. Percentages of overlap was determined by the following formula: (number of loci in a particular category overlapping genetic elements) / (total number of loci in that category)* 100. Predominant read size and genomic strand polarity of all loci was determined by the initial run of merged wild-type and *rdr1/2/6* datasets.

In order to determine the number of reads aligning to different portions of overlapping genes, sequences corresponding to all of *A. thaliana* TAIR10 gene models were extracted from the *A. thaliana* TAIR10 genome using GetFasta from the Bedtools suite (Quinlan, 2014). Our wild-type libraries were aligned against the gene sequences using ShortStack 3.8.1. The start and stop sites of the untranslated regions, exons, and introns relative to gene sequences was calculated using the coordinates provided by The Arabidopsis Information Resource (Rhee et al., 2004) and the amount of reads in each feature of the genes was determine using CoverageBed of the Bedtools suite (Quinlan, 2014). The observed number of reads in each feature was divided by length of that feature in order to mitigate sequencing biases.

### Expression analysis of unclassified small RNAs in various mutant libraries

Small RNA libraries corresponding to several different types of DCL mutants were downloaded from the Gene Expression Omnibus. A list of these libraries is shown in Table S1. In each case, the mutant and corresponding wild-type libraries were trimmed and aligned against the *A. thaliana* genomes using ShortStack ver 3.8.1 using the results file from our initial run as a ‘locifile.’ Differentially expression analysis was performed as before using DESeq2 (Love et al. 2014), except no global miRNA-based normalization was used for *rrpl1*, *rrpl2*, *rtl2*, *dcl1*, *dcl234*, and *dcl3* mutant libraries.

### Conservation analysis of different types of loci

Conservation data for each nucleotide in the *A. thaliana* TAIR10 genome was from Haudry et al. 2013. Briefly, genomes from nine distinct crucifer species (*A. thaliana, Leavenworthia alabamica, Sisymbrium irio, Aethionema arabicum, Capsella rubella, A. lyrata, Brassica rapa, Eutrema salsugineum*, and *Schrenkiella parvula*) were aligned in a pairwise fashion. Using *A. thaliana* (TAIR10) as a reference, the conservation of each nucleotide was assigned values in arbitrary units based on the selection pressures on that nucleotide (Haudry et al. 2013).

An in-house Python script (available upon request) was used to compare the conservation level of a nucleotide in each locus described in this study. The conservation level of each nucleotide was added to a list recurrent with the depth of coverage of that nucleotide in merged wild-type alignments. Finally, the median of conservation data of the locus was then calculated.

### Alignment and expression of *A. thaliana* mRNAs

Two libraries corresponding to *A. thaliana* transcriptomic data in wild-type inflorescence tissues (Table S1) were aligned against the TAIR10 genome using Tophat2 (Kim et al., 2013). The aforementioned list of TAIR10 gene models was used as the gene query file in the Tophat2 analysis.

## Author Contributions

MJA conceived of the project. SP performed all experiments and analyses. MJA and SP wrote the manuscript.

## Acknowledgments

We thank Michael Schon and Michael Nodine for the conservation data. We also thank Dr. Naomi Altman for helpful comments and advice concerning the statistics of the DESeq2 package. Ceyda Coruh prepared small RNA libraries. The authors thank Penn State University Genomic Core Facility for small RNA-seq sequencing, and all members of the Axtell Lab for constructive comments.

## Short Supporting Information Legends

**Figure S1.** The *MIR5635* paralogs are *RDR1/2/6*-dependent and overlap with long, inverted repeats.

**Figure S2.** Examples of different *rdr1/2/6*-independent categories.

**Figure S3.** Relative accumulation and differential expression of unclassified *RDR1/2/6*-independent clusters in various sRNA mutants.

**Figure S4.** Examples of gene-intersecting, unclassified *RDR1/2/6*-independent loci with different relative polarities.

**Table S1.** RNA libraries used in this study.

**Dataset S1.** Many miRBase21-collated loci are rdr1/2/6-dependent and are not *MIRNAs*.

**Dataset S2.** *de novo Small RNA clusters annotated in this study*.

**Dataset S3.** Differential expression status of *RDR2*-dependent and RDR6-dependent loci.

**Dataset S4.** Differential expression status of known *rdr1/2/6*-independent clusters.

**Dataset S5.** Ontologies of unclassified sRNA clusters-intersecting genes divided by their relative polarities.

---

**Figure S1.**
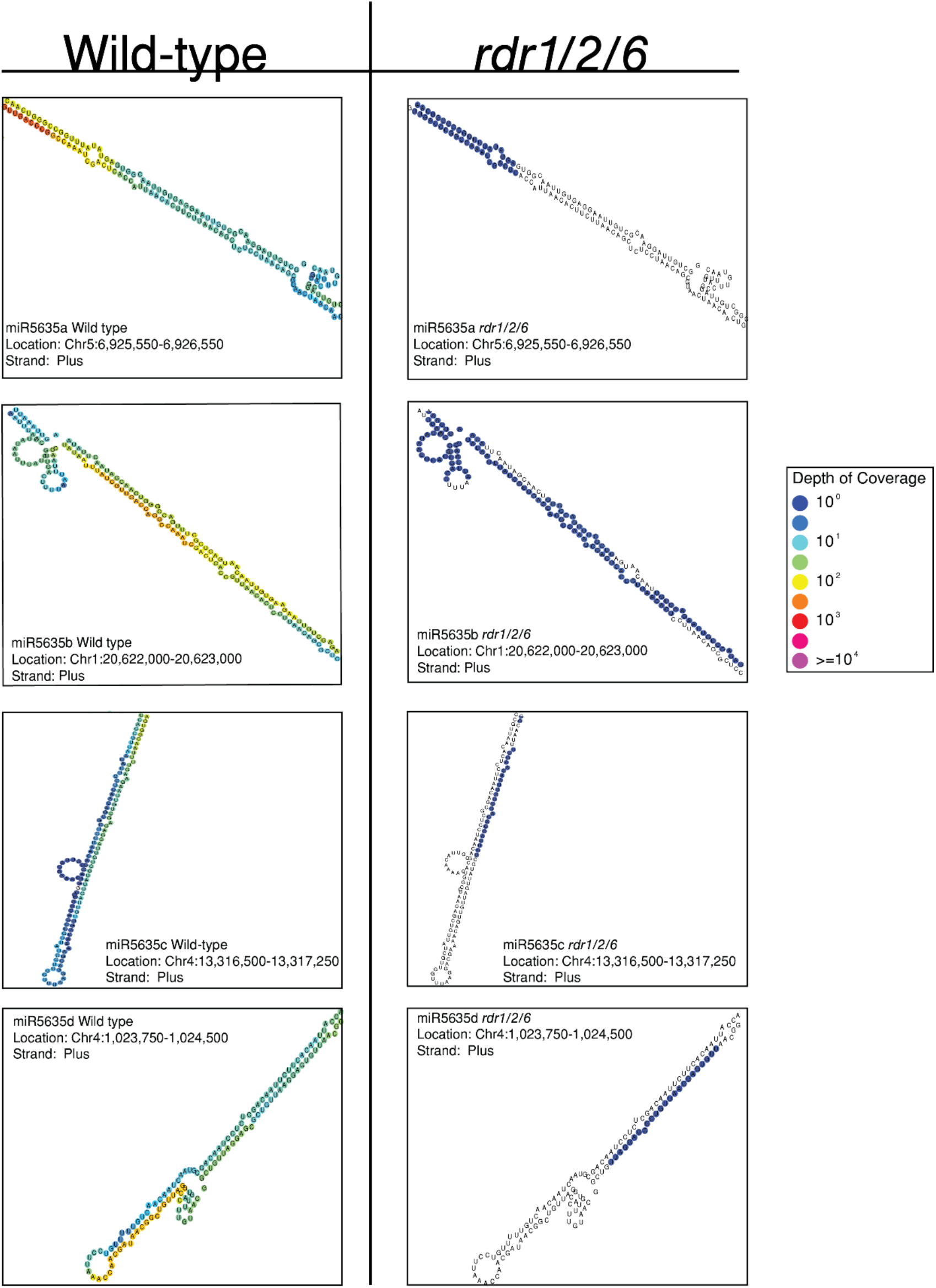
The *MIR5635* paralogs are *RDR1/2/6*-dependent and overlap with long, inverted repeats. Part of the predicted secondary structures of the RNA with coverage of each nucleotide is shown is shown using merged wild type and *rdr1/2/6* libraries. Colors show the depth of coverage as indicated by the legend. No color means no reads mapped to the particular nucleotide

**Figure S2.**
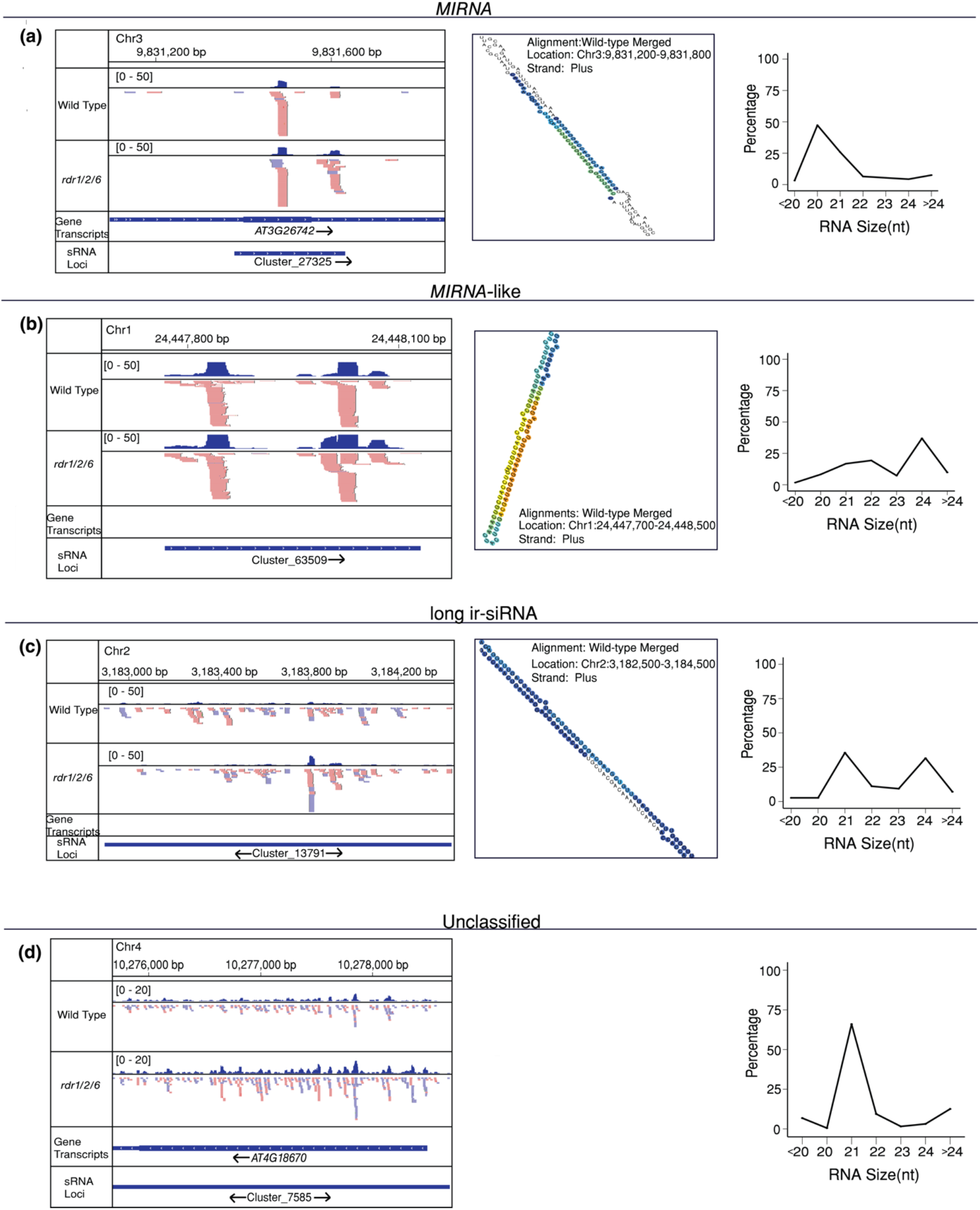
Examples of different *rdr1/2/6*-independent categories. (a) Alignments, secondary structure, and read size distribution of an example of a *MIRNA*. Arrows represent genomic polarity of locus. Two arrows mean that the particular locus is derived from both strands.
(b) Same as panel (a) but for an example of a *MIRNA*-like locus.
(c) Same as panel (a) but for an example of a long ir-siRNA locus.
(d) Same as panel (a) but for an example of an unclassified locus.

**Figure S3.**
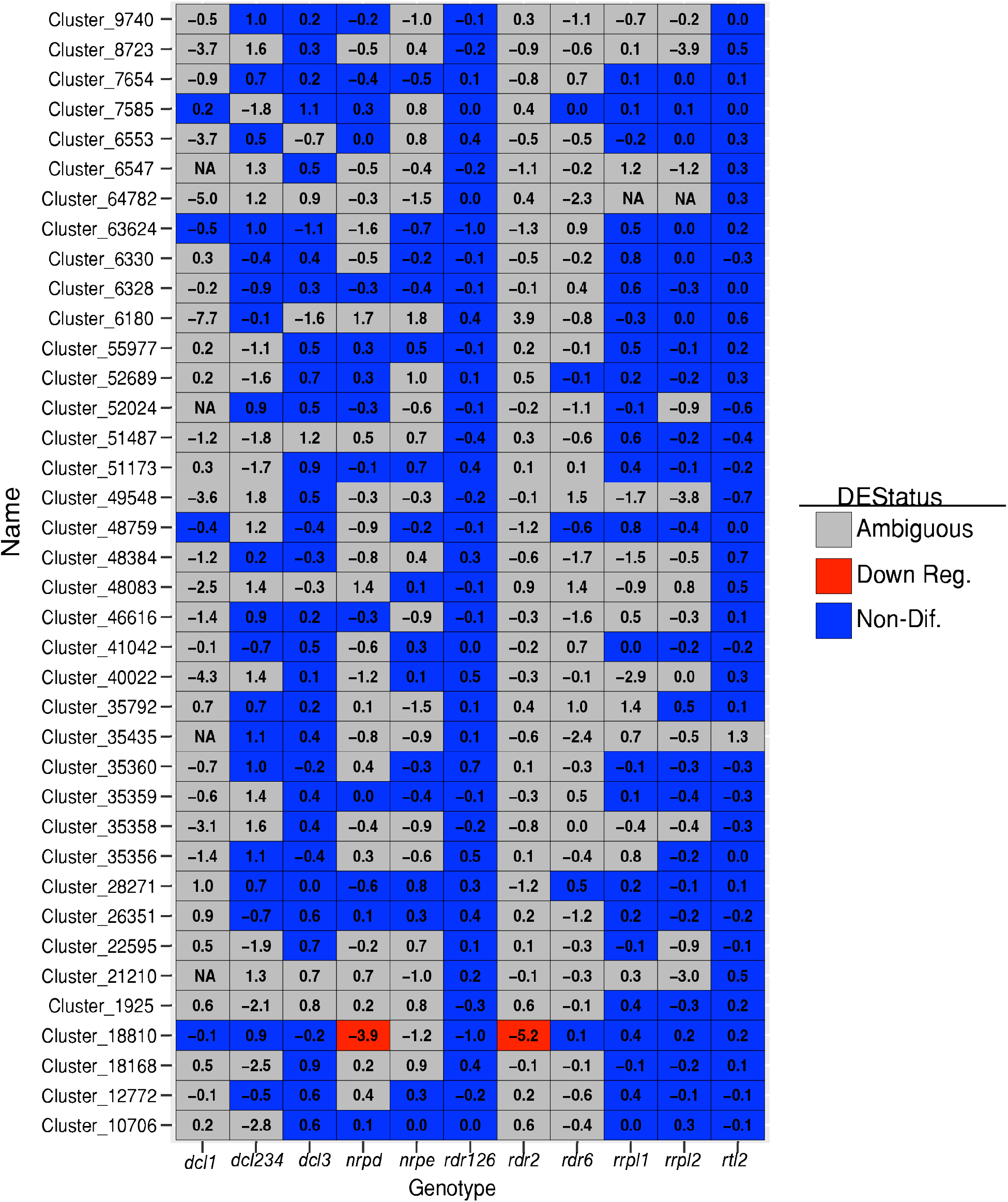
Relative accumulation and differential expression of unclassified *RDR1/2/6*-independent clusters in various sRNA mutants. Heatmap showing normalized sRNA accumulation levels of unclassified loci in different mutant libraries relative to wild-type. The ratio of small RNA accumulation in the indicated genotypes over that in corresponding wild-type library was computed via DESeq2. Table S1 shows details of the sRNA-seq libraries.

**Figure S4.**
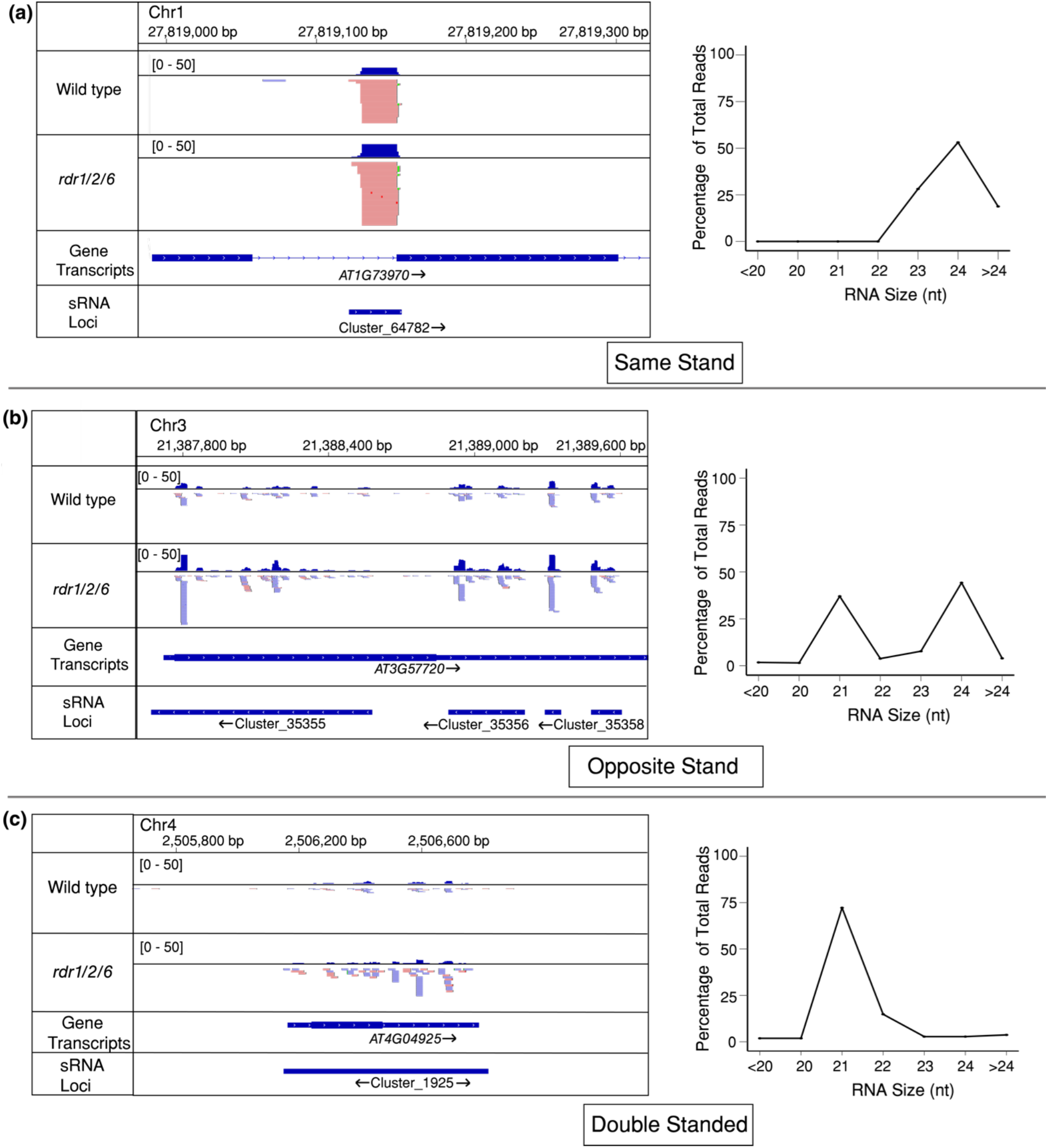
Examples of gene-intersecting, unclassified *RDR1/2/6*-independent loci with different relative polarities. (a) Alignments and read size distribution of an example of an unclassified sRNA locus with the same polarity as the gene it intersects. Red and blue reads align to positive and negative strands, respectively. Arrows represent genomic polarity of locus. Two arrows mean that the particular locus is derived from both strands.
(b) Same as panel (a) except for an unclassified sRNA locus that has the opposite polarity as the gene it intersects.
(c) Same as panel (a) except for an unclassified sRNA locus which is derived from both strands of the genome.

**Table S1.**
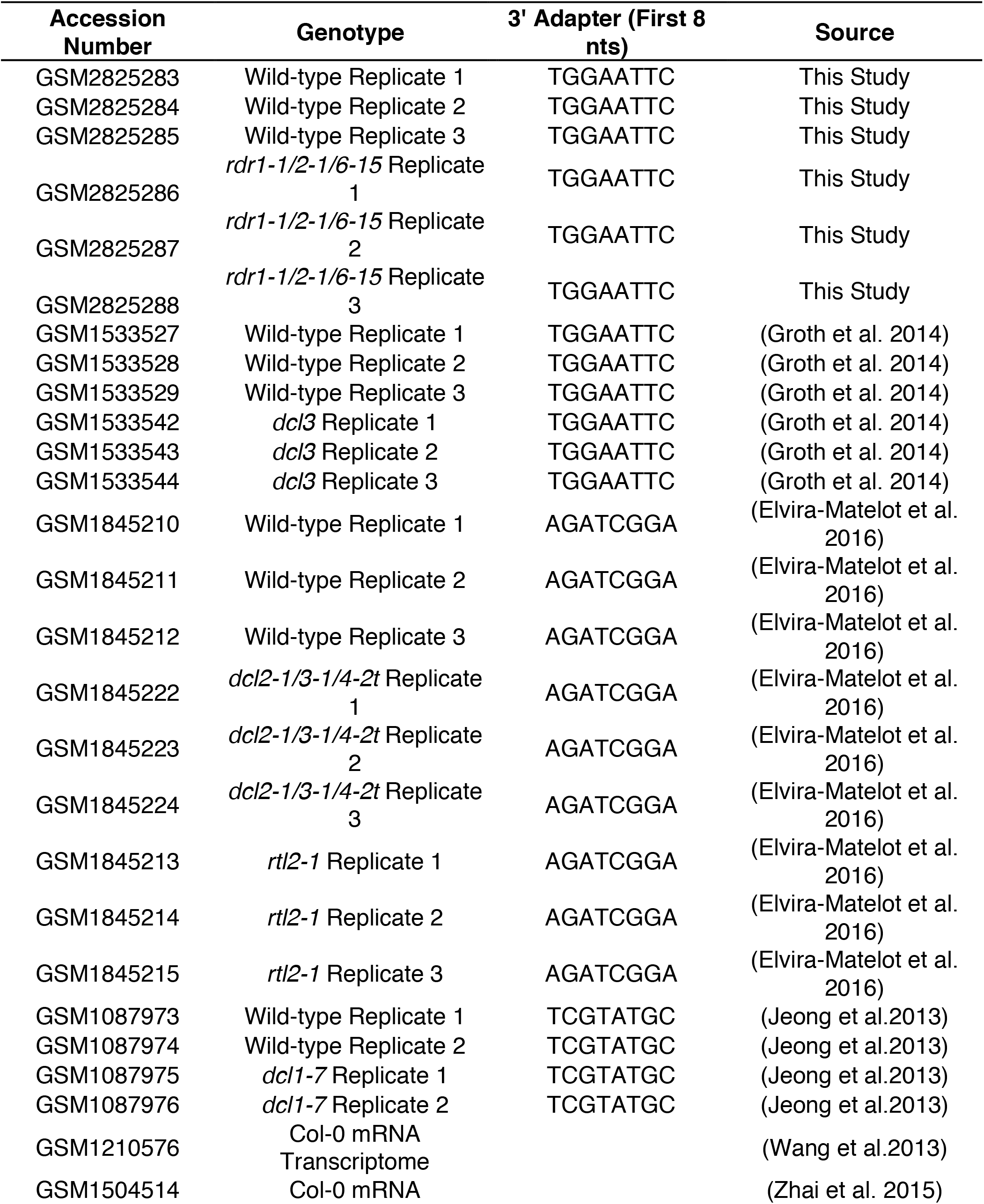
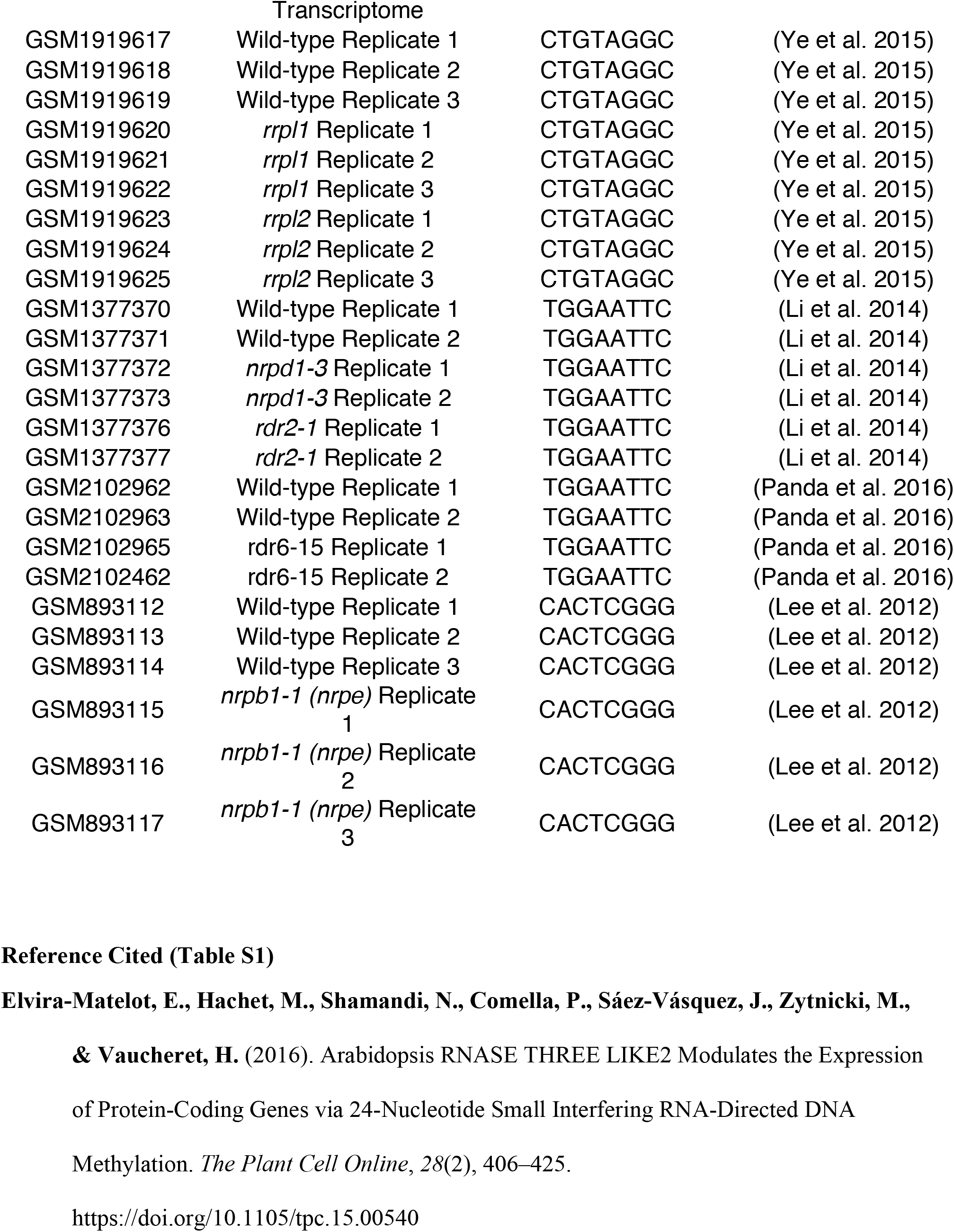
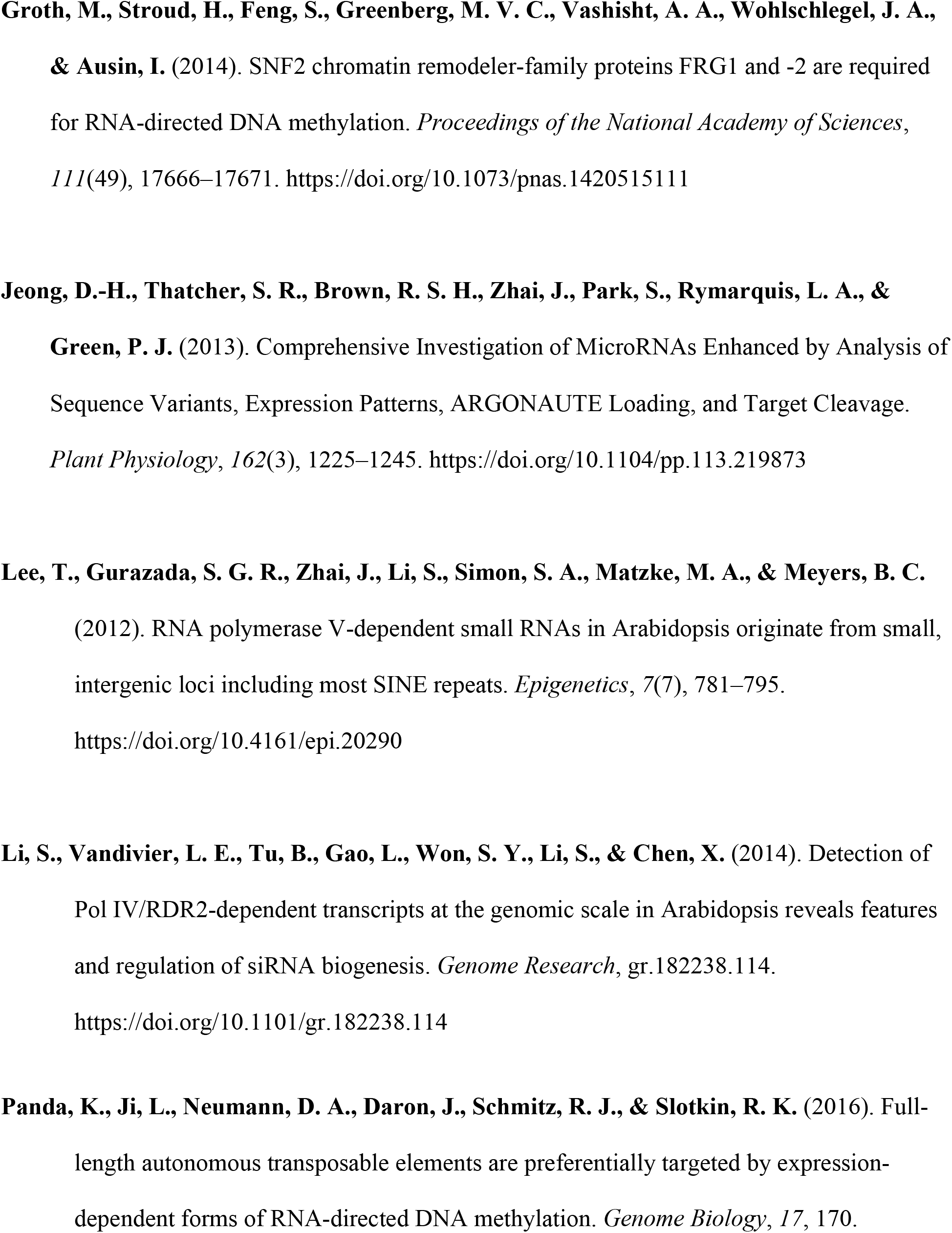
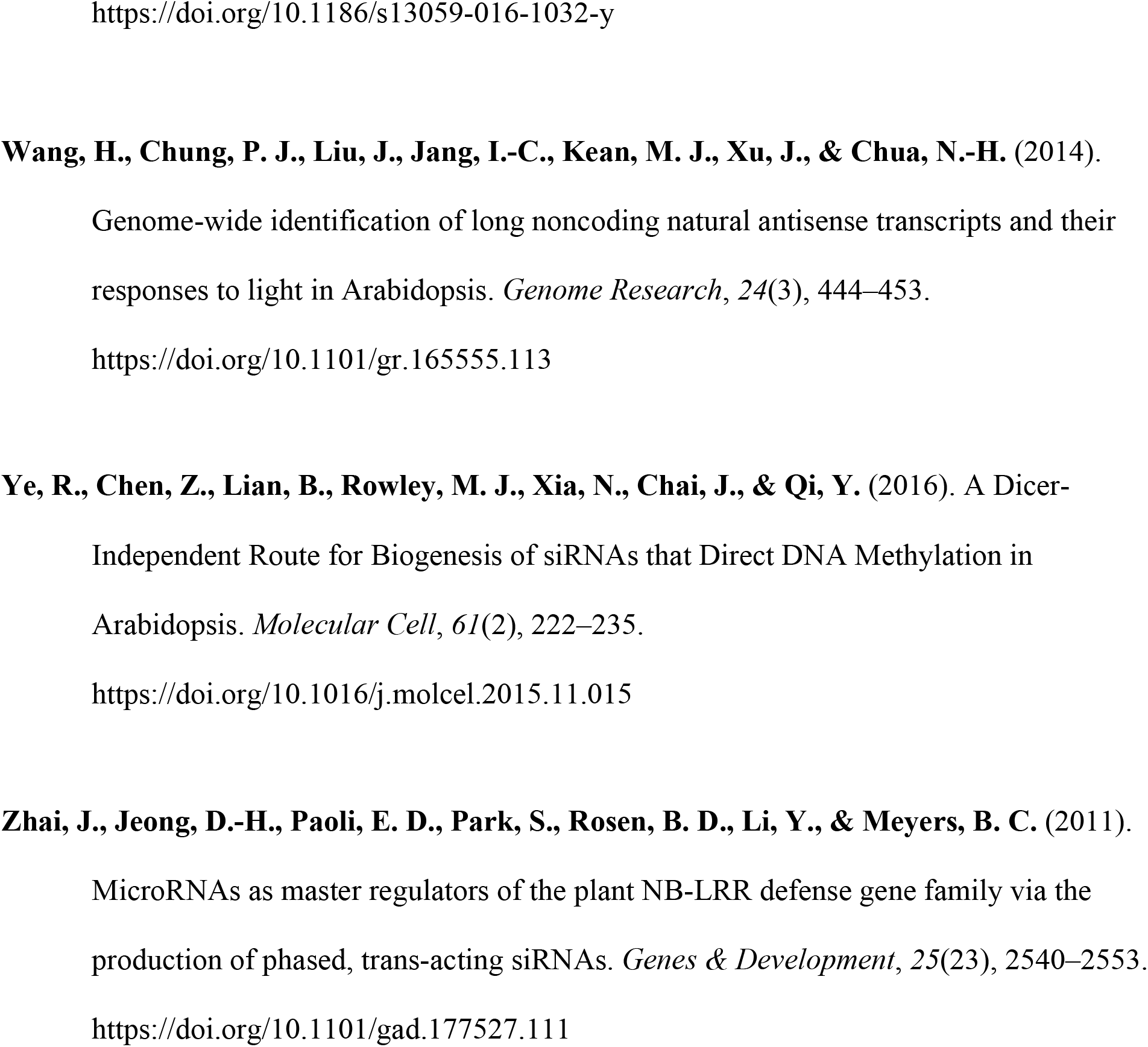
RNA libraries used in this study.

